# N1-Methylpseudouridine and pseudouridine modifications modulate mRNA decoding during translation

**DOI:** 10.1101/2022.06.13.495988

**Authors:** Jeremy G. Monroe, Lili Mitchell, Indrajit Deb, Bijoyita Roy, Aaron T. Frank, Kristin Koutmou

**Affiliations:** Department of Chemistry, University of Michigan, Ann Arbor, MI 48109; RNA and Genome Editing, New England Biolabs Inc., Ipswich, MA 01938; Department of Biophysics, University of Michigan, Ann Arbor, MI 48109

**Keywords:** N1-methylpseudouridine, ribosome, RNA modification, translation, miscoding

## Abstract

The ribosome relies on hydrogen bonding interactions between mRNA codons and incoming aminoacyl-tRNAs to ensure rapid and accurate protein production. The inclusion of chemically modified bases into mRNAs has the potential to alter the strength and pattern of hydrogen bonding between mRNAs and aminoacyl-tRNAs to alter protein synthesis. We investigated how the N1-methylpseudouridine (m^1^Ψ) modification, commonly incorporated into therapeutic and vaccine mRNA sequences, influences the ability of codons to react with cognate and near-cognate tRNAs and release factors. We find that the presence of a single m^1^Ψ does not substantially change the rate constants for amino acid addition by cognate tRNAs or termination by release factors. However, insertion of m^1^Ψ can affect the selection of near-cognate tRNAs both *in vitro* and in human cells. Our observations demonstrate that m^1^Ψ, and the related naturally occurring pseudouridine (Ψ) modification, exhibit the ability to both increase and decrease the extent of amino acid misincorporation in a codon-position and tRNA dependent manner. To ascertain the chemical logic for our biochemical and cellular observations, we computationally modeled tRNA^Ile(GAU)^ bound to unmodified and m^1^Ψ- or Ψ-modified phenylalanine codons (UUU). Our modeling suggests that changes in the energetics of mRNA:tRNA interactions largely correlate with the context specificity of Ile-miscoding events we observe on Ψ and m^1^Ψ containing Phe codons. These studies reveal that the sequence context of a given modification within an mRNA plays a large role in determining how (and if) the modification impacts the number and distribution of proteoforms synthesized by the ribosome.

## INTRODUCTION

Chemically modified nucleosides are present in all organisms, often playing essential roles in key cellular processes including splicing and translation (1–3). Defects in ribosomal RNA (rRNA) and transfer RNA (tRNA) modifying enzymes are linked to a host of deleterious human health outcomes, illustrating the importance of RNA modifications in protein synthesis (4–7). There are over 150 unique modifications reported in RNAs that range in size and complexity from isomerized or saturated nucleosides (*e.g*. pseudouridine and dihydrouridine) to large chemically diverse functional groups (*e.g*. NAD^+^, N(6)-threonylcarbamoyladen-osine, glycan and farnesyl) (8–11). RNA modifications have been widely studied for almost three quarters of a century and until recently were thought to be almost exclusively incorporated into non-coding RNA species (ncRNAs). However, the transcriptome wide mapping of 13 RNA modifications revealed that protein coding messenger RNAs (mRNAs) can also contain modifications at thousands of sites (12–26). This discovery has raised the possibility that mRNA modifications might play a previously underappreciated role in post-transcriptionally regulating gene expression.

The majority of enzymes that modify mRNAs also catalyze their incorporation into ncRNAs central to protein synthesis (8). Like their protein post-translational counterparts, mRNA post-transcriptional modifications are generally present at sub-stoichiometric levels, with transcripts existing in a mixed population of modified and unmodified states (15, 26–28). Together these circumstances make ascertaining the impact of mRNA modifications on translation challenging. In cells, any changes to protein output observed when RNA modifying enzymes are removed are difficult to directly attribute to alterations in a particular mRNA’s modification status. Therefore, studies using reconstituted translation systems, where it is possible to uniformly change the modification status of an mRNA without impacting that of ncRNA, have been particularly useful for assessing the consequences of mRNA modifications on translation (29). Initial studies reveal that modifications commonly slow the ribosome, though some only do so only in particular mRNA sequence contexts (29). Additionally, a subset of mRNA modifications, including pseudouridine (Ψ) and inosine (I), also impact the accuracy of mRNA decoding (30–32). These findings suggest that there is a broad range of possible consequences when the ribosome encounters an mRNA modification. Developing a framework for understanding how individual modifications impact translation in differing sequence contexts will be crucial as researchers seek to uncover which of the thousands of chemically modified positions reported in mRNA codons are the most likely to have consequences for protein synthesis in cells.

In addition to being present in naturally occurring RNA molecules, modifications are also heavily incorporated in RNA-based therapeutics and mRNA vaccines (33–36). Indeed, the mRNA transcripts that form the basis of many currently available COVID-19 mRNA vaccines substitute every uridine nucleoside with N1-methylpseudouridine (m^1^Ψ) (37). The addition of m^1^Ψ limits the cellular innate immune response to dramatically stabilize the mRNA transcript, and ultimately increase the amount of protein synthesized (38–41). Recent studies in a lysate-based translation system suggest that m^1^Ψ slows the ribosome in a manner that can be alleviated by the addition of membranes (42). However, there is limited information available directly measuring how m^1^Ψ influences the rate and accuracy of amino acid addition. This is an important question to ask because m^1^Ψ shares much of its structure with pseudouridine (Ψ) (Figure 1A), a modification that has been shown to change translation speed and tRNA selection (30, 43, 44). Even subtle changes in translation rates or fidelity have the potential to impact protein folding or function (45–48). Therefore, establishing if there are situations in which m^1^Ψ can alter translation will be critical for the continued development of mRNA-based therapeutics and vaccines in addition to understanding how different types of chemical moieties impact translation.

**Figure 1.**
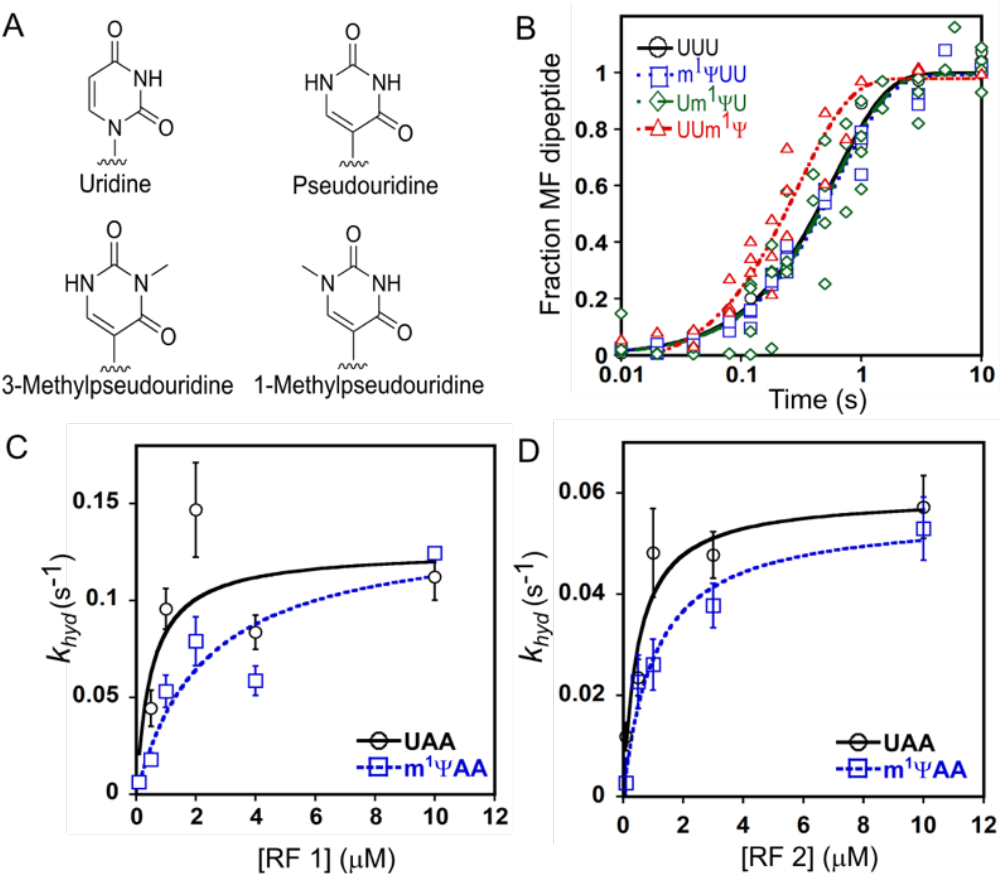
Cognate amino acid addition is modestly increased on UUm^1^Ψ, but not m^1^ΨUU or UUm^1^YU, codons. (**A**) The chemical structures of the nucleobases we investigated. (**B**) The formation of Met-Phe (MF) dipeptide as a function of time by *E. coli* ribosomes containing ^35^S-^f^Met-tRNA^fMet^ bound to an AUG start codon in the P site, and unmodified (◯-UUU) or modified (□-m^1^ΨUU, ◇-Um^1^ΨU, □-UUm^1^Ψ) codons in the A site. (**C**) The *K*_½_ curve for RF1. *k*_obs_ values for RF1-catalyzed 35S-^f^Met release on UAA (◯) or m^1^ΨAA (□) are displayed as a function of [RF1]. (**D**) The *K*_½_ curve for RF2. *k*_obs_ values for RF2-catalyzed ^35^S-^f^Met release on an UAA (◯)or m^1^ΨAA (□) are displayed as a function of [RF2].

To ascertain the molecular level consequences of m^1^Ψ codon modifications on ribosome decoding, we compared the translation of unmodified and nPΨ-modified codons in both a fully reconstituted bacterial *in vitro* translation system and HEK293 cells. These studies reveal that in contrast to Ψ, the presence of a single m^1^Ψ does not reduce the rate constant for cognate amino acid addition. However, m^1^Ψ does influence the accuracy of amino acid addition. We demonstrate that Ψ and m^1^Ψ can both impede and enhance near-cognate tRNA selection depending on the surrounding sequence context and identity of the tRNA. Comparison how Ψ and m^1^Ψ modifications affect translation reveal that uridine base isomerization and methylation each contribute to the ability of m^1^Ψ to perturb mRNA decoding. Computational modeling of a near cognate tRNA^Ile,UAG^ bound to modified and unmodified Phe (UUU) codons in the context of the A site suggests that changes in the energies of mRNA:tRNA interactions likely account for the context dependent effects of Ψ and m^1^Ψ we observe. These findings demonstrate that Ψ and m^1^ Ψ modulate ribosome decoding and have the potential to modulate the speed and accuracy of protein production from both native and therapeutic mRNA sequences.

## RESULTS

### m^1^Ψ modestly impacts the rate constant for Phe addition and K_½_ for peptide release

We used a fully reconstituted *E. coli in vitro* translation system to evaluate the consequences of incorporating m^1^Ψ into mRNA codons on translation elongation and termination. In contrast to reporter-based studies in cells and lysates, the *in vitro* system we implemented is not influenced by extra-translational factors that can change observed protein levels (*e.g*. RNases and proteases) and allows us to directly examine individual steps along the translation pathway with high resolution (49). This *E. coli* translation system has long been used to study translation elongation because tRNA binding sites and ribosome peptidyl-transfer center are highly conserved between eukaryotic and bacterial ribosomes (50, 51).

The rate constants for amino acid addition were measured on unmodified (UUU) and m^1^Ψ modified (m^1^ΨUU, Um^1^ΨU, and UUm^1^Ψ) phenylalanine (Phe) codons (Figure 1B). We chose to first evaluate amino acid incorporation rates on a UUU codon because the kinetics of Phe addition on UUU is well established, and UUU codons are present in the Pfizer/BioNTech mRNA COVID-19 vaccine (37). Our translation reactions were initiated by mixing *E. coli* 70S ribosome initiation complexes (ICs; ^35^S-labeled formylmethi-onine-tRNA^fMet^ [^35^S-^f^Met] bound to an AUG in the P site and Phe codon in the A site) with an excess of ternary complexes (TCs; Phe-tRNA^Phe^•EF-Tu•GTP). Reactions were quenched at select time points, and the unreacted ^35^S-^f^Met and ^35^S-^f^Met-Phe products are visualized by electrophoretic TLC (eTLC) (SI Figure 1A). These studies reveal that cognate Phe incorporation on m^1^Ψ modified codons is largely unchanged, though we observe a slight (2 ± 0.3-fold) increase in the rate constant for Phe addition when m^1^Ψ is in the third position in the codon (Figure 1B, S1 Figure 1B, SI Table 1).

All three stop codons begin with uridine (UAA, UAG, UGA) ensuring that modified stop codons will be present in synthetic mRNA-based vaccines and therapeutics. We evaluated the ability of bacterial class I release factors (RF1 and RF2) to hydrolyze peptidyl-tRNA bonds and terminate translation on m^1^Ψ modified stop codons. To accomplish this, we reacted termination complexes (*E. coli* 70S ribosomes with ^35^S-^f^Met bound to an AUG in the P site, and a universal stop codon positioned in the A site (UAA, m^1^ΨAA)) with varying concentrations of RF1 and RF2 (0.1-10 μM). The reactions were quenched at a range of time points and ^35^S-^f^Met hydrolyzed by RFs was detected on an eTLC (S1 Figures 2 and 3). At saturating levels of RF1 and RF2 the rate constants for peptide release (*k_hyd,max_*) on UAA and m^1^ΨAA codons are equivalent (~ 0.1 s^-1^) and comparable previously published termination rates on an unmodified UAA codon (S1 Tables 2 and 3) (52, 53). However, this was not the case at sub-saturating conditions, as reflected by the 2- to 4-fold increase in *K*_½_ values obtained for peptidyl-tRNA hydrolysis by RF1 and RF2 on a m^1^ΨAA (Figure 1C-D, SI Tables 2 and 3). The redistribution of the electronegativity around the pyrimidine ring in m^1^Ψ can weaken the hydrogen bonding network between the first two nucleotides of the stop codon and release factors, perhaps accounting for this observation (53, 54). Nonetheless, because *k_hyd,max_* is unperturbed we do not expect m^1^Ψ to impede translation termination in cells unless the concentration of release factors becomes severely limited, or, the termination codon is in a particularly poor sequence context (55). This supposition is supported by numerous observations that reporter peptides and therapeutic RNA sequences generated from fully m^1^Ψ-substituted mRNAs yield protein products of the expected length (38, 56).

### m^1^Ψ influences aminoacyl-tRNA selection by the ribosome in a context dependent manner

Chemical modifications to nucleobases can change the propensity of the ribosome to incorporate alternative amino acids into a growing polypeptide chain (29–32, 57–60). In comparison to uridine, m^1^Ψ possesses a repositioned, methylated nitrogen in its pyrimidine ring (Figure 1A). These two changes provide m^1^Ψ the opportunity to alter the conformational fit of an mRNA in the ribosome, and increase the variety of mRNA:tRNA base pairing interactions possible. Consistent with this idea, Ψ, which shares a repositioned nitrogen with m^1^Ψ, was previously shown to enhance the reaction of some near-cognate tRNAs on UUU codons *in vitro* and in HEK293 cells (30). To begin examining if m^1^Ψ similarly influences aminoacyl-tRNA (aa-tRNA) selection, we qualitatively evaluated the impact of m^1^Ψ on the propensity of Phe codons to react with near-cognate tRNAs. In these assays, 70S *E. coli* initiation complexes were generated with unmodified (UUU) and modified (m^1^ΨUU, Um^1^ΨU, and UUm^1^Ψ) codons in the A site, and reacted with EF-Tu containing ternary complexes formed using a mixture of total tRNA aminoacylated either by reacting all 20 amino acids with S100 (total aa-tRNA^aa^), or with a single amino acid and aminoacyl tRNA synthetase (Phe-tRNA^Phe^, Ser-tRNA^Ser^, Leu-tRNA^L^e^u^, Ile-tRNA^Il^e and Val-tRNA^val^) (49). Relative to UUU, we observed that m^1^ΨUU appears to react more robustly with total aa-tRNA^aa^, and promote the production of higher-levels of miscoded Met-Ile (MI) and Met-Met-Val (MV) peptides (Figure 2A). Um^1^ΨU and UUm^1^Ψ codons also exhibit different levels of reactivity with near-cognate tRNAs than an unmodified UUU (S1 Figure 4).

**Figure 2.**
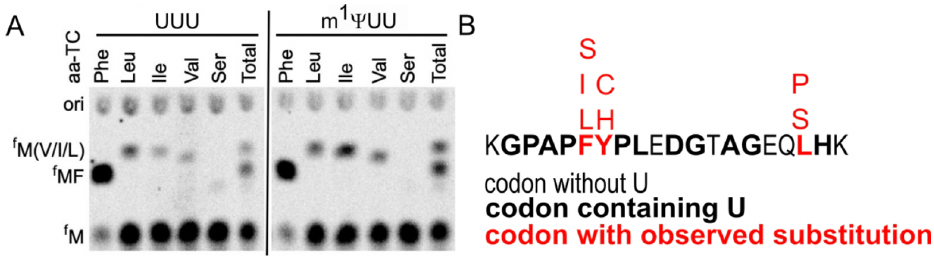
m^1^Ψ amino acid selectivity *in vitro* and in HEK293 cells. **(A)** eTLC displaying dipeptide products from translation reactions performed with 70S initiation complexes (ICs) with an unmodified UUU or m^1^ΨUU codon in the A site and total *E. coli* tRNA aminoacylated with a single amino acid (aa-TC). Relative to ICs containing a UUU codon in the A site, higher levels of miscoded MI and MV dipeptide products and lower levels of MS were generated from m^1^ΨUU containing ICs. **(B)** Summary of amino acid substitutions observed by mass spectrometry in a luciferase peptide incorporated on m^1^Ψ-containing mRNAs translated in HEK293 cells (SI Table 8).

Using the information obtained from our qualitative screens, we selected tRNAs for closer, quantitative investigation. We choose to evaluate three near-cognate tRNAs that either appeared to react more, less, or to the same extent on unmodified and m^1^Ψ modified codons (tRNA^Ile(GAU)^, tRNA^Leu(CAG)^ and tRNA^Ser(UGA)^) (Figure 2A). The single-turnover rate constants (*k*_obs_) for each amino acid mis-incorpo-rating on unmodified and m^1^Ψ substituted Phe (UUU) codons was measured at saturating concentrations of individually purified charged aa-tRNA (Figure 3, S1 Figures 5 and 6). An energy regeneration mix was included in our reactions to permit the reformation of ternary complexes (aa-tRNA:EF-Tu:GTP) after an aa-tRNA is rejected by the ribosome (61). We find that tRNA identity and the position of m^1^Ψ within a codon influence the rate constants for amino acid substitution (Figure 3, S1 Table 4). For example, m^1^Ψ substitution at the first position in the Phe codon (m^1^ΨUU) does not change the *k*_obs_ values for Leu or Ser incorporation, but increases the rate constant for Ile addition by 2.2 ± 0.4-fold (Figures 3D and 4A, SI Table 4). This differs markedly from what we observed on Um^1^ΨU-modified codons, which have a much larger effect on tRNA selection. The *k*_obs_ values are significantly reduced for Ile (10 ± 2-fold) and Leu (4 ± 1-fold) addition, while the rate constant for Ser misincorporation is conversely increased by 3.5 ± 0.4-fold (Figure 3D, SI Table 4). Substitution at the wobble position (UUm^1^Ψ) generally had modest impacts on the rate constants for amino acid incorporation; decreasing the *k*_obs_ for Leu addition (2 ± 0.3-fold), while not impacting the *k*_obs_ values for Ile and Ser addition (Figure 3D, SI Table 4). The findings of our kinetic studies are generally consistent with our initial qualitative assays (Figures 2, 3 and S1 Figure 5), and together indicate that m^1^Ψ codon modifications can both increase and decrease the ability of Phe UUU codons to react with near-cognate tRNAs in the A site.

**Figure 3.**
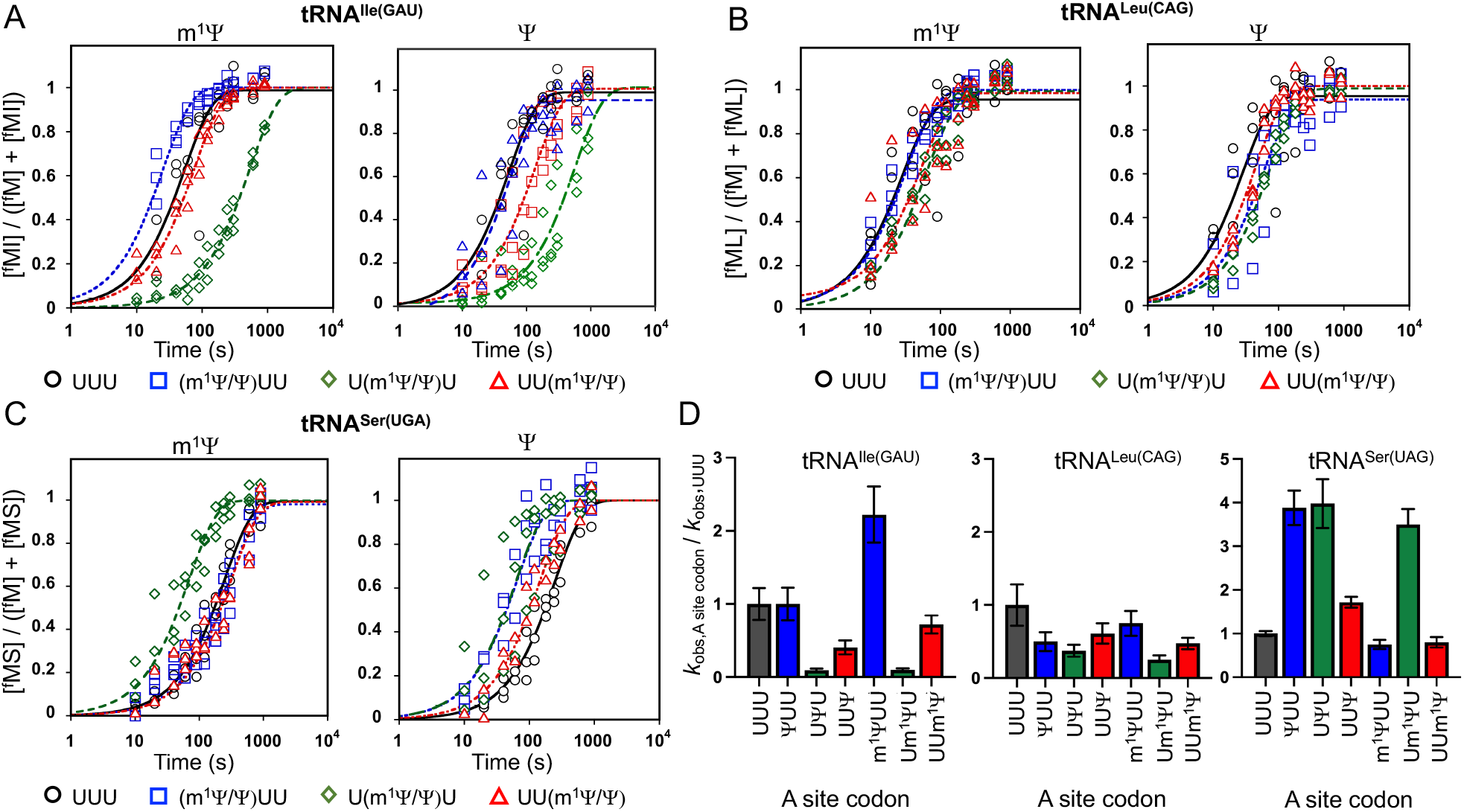
Ψ and m^1^Ψ impact the rates of the ribosome reacting with near-cognate tRNAs in a sequence context dependent manner. Plots of dipeptide formation as a function of time (seconds). Miscoding reactions were performed with *E. coli* ICs containing 3≡S-fMet-tRNAfMθt bound to an AUG start codon in the P site, and unmodified (◯-UUU) or modified (□-Ψ/m^1^ΨUU, ◇ - UΨ/m^1^ΨU, □-UUΨ/m^1^Ψ) codons in the A site. Purified ICs were reacted with TCs containing **(A)** Ile-tRNA^Ile(GAU)^, Leu-tRNA^Leu(CAG)^), or **(C)** Ser-tRNA^Ser(UGA)^. **(D)** The rate constants (*k*_obs_) for isoleucine, leucine, and serine misincorporation on Ψ- and m^1^Ψ-modified codons relative to the rate constants for isoleucine, leucine, and serine misincorporation on a UUU codon.

**Figure 4.**
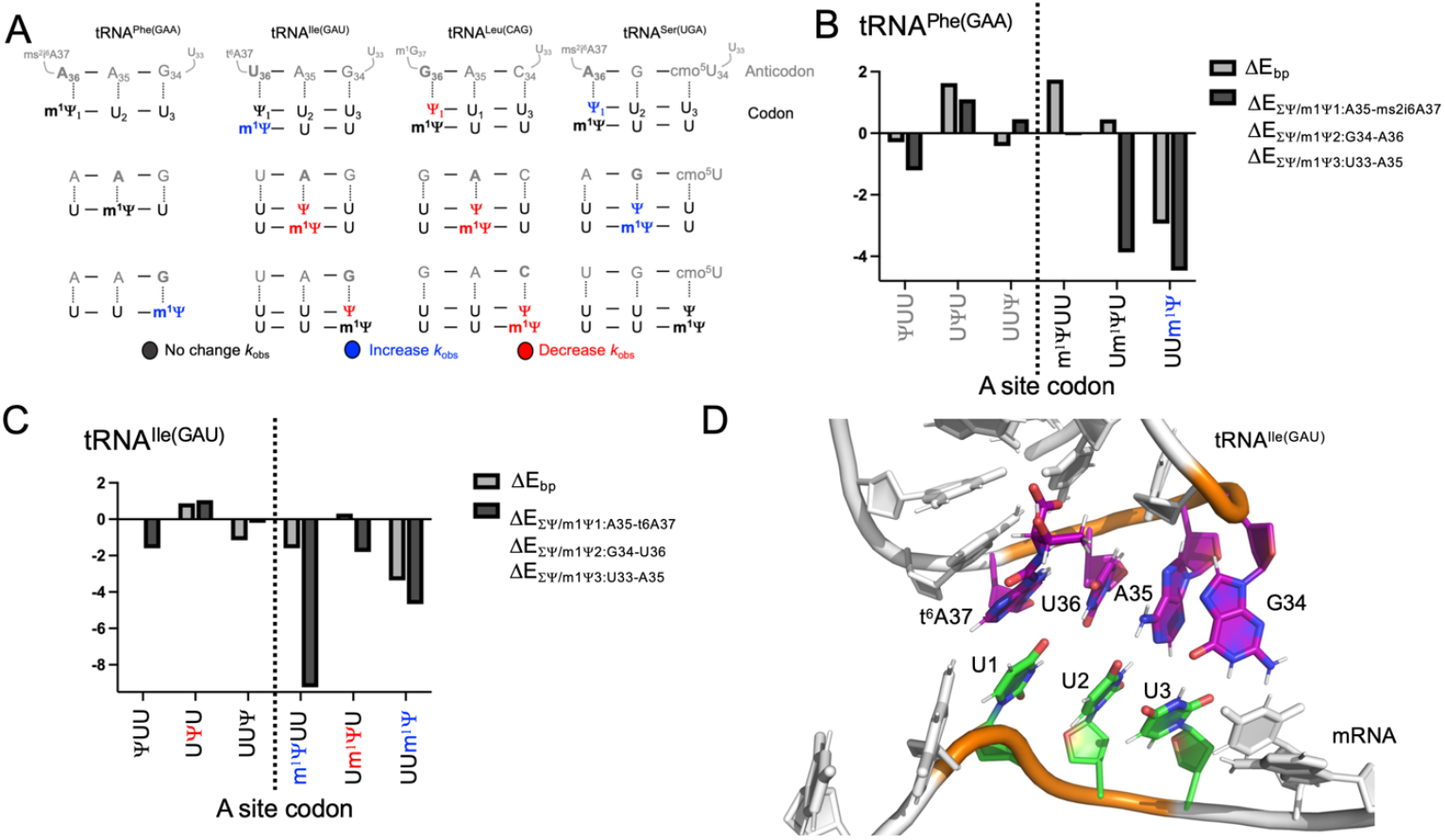
Changes in the energetics of mRNA:tRNA interactions correlate with observed differences in Phe and Ile in-corporation on Ψ - and m Ψ-containing codons. **(A)** Summary of data in SI Tables 4 and 5 displaying how a Ψ and m^1^Ψ impact the rate constants for the reaction of near-cognate tRNAs. (**B and C)** Summary of MM data. Gray bars reflect the change in energy for interactions between a modified mRNA position (Ψ/m^1^Ψ) and the base paring (**B**) tRNA^Phe(GAA)^ or (**C**) tRNA^Ile(GAU)^ residue (ΔEbp) relative to an unmodified mRNA U. Black bars reflect the total change in energy (ΔE_ΣΨ/m1Ψ:X-Y_) for interactions between a modified mRNA position (Ψ/m^1^Ψ) and three **(B)** tRNA^Phe(GAA)^ or **(C)** tRNA^Ile(GAU)^ residues (the base paired nucleotide, and nucleotides 5’ and 3’ the bp) relative to an unmodified mRNA U. **(D)** Molecular model of an unmodified sequence coding for a Phe UUU codon and a tRNA^Ile(GAU)^. The hypermodification t^6^A37 is also shown the tRNA.

### Uridine isomerization contributes to observed changes in amino acid substitution on m^1^Ψ containing codons

To determine how the C5-glycoside uridine isomerization and an N1-methylation individually impact the ability of m^1^Ψ to modulate amino acid misincorporation, we measured the rate constants for Leu, Ile and Ser addition on Ψ modified Phe codons (ΨUU, UΨU and UUΨ). Ψ was selected for study because it contains the same isomerized uridine base as m^1^Ψ, but lacks the methylation an position N1 (Figure 1A). The impact of Ψ on Ile and Leu insertion was similar to what we observed when m^1^Ψ is present in codons (Figure 3, S1 Figure 7). For example, the rate constant for Ile is significantly decreased (10 ± 2-fold) when Ψ is incorporated at the second codon position (UΨU), while Leu is added more slowly when Ψ is at all three positions in the codon (Figure 3, S1 Table 5). In contrast to m^1^Ψ, Ser incorporation occurs with a 4 ± 0.5-fold faster rate constant when Ψ is at the first and second positions in the codon, and is not influenced by Ψ substitution at the wobble position (UUΨ). These observations indicate that uridine isomerization largely accounts for changes in how the ribosome decode some tRNAs on for m^1^Ψ-substituted mRNAs. However, comparison of the reactivity profiles Ψ and m^1^ΨUU reveal the N1 methyl group can suppress the effect of uridine isomerization on the rate constants for amino acid misincorporation by other near-cognate tRNAs (*e.g*. ΨUU vs m^1^ΨUU reacting with tRNA^Ser(UGA)^) (Figure 3D, 4A).

### Amino acid substitution in HEK293 cells increases on some m^1^Ψ containing codons

Our *in vitro* translation data reveal that m^1^Ψ and Ψ affect tRNA selection by *E. coli* ribosomes in different ways depending on their sequence context. We next asked if m^1^Ψ has similar effects on amino acid selection in eukaryotic cells. To approach this question, we transfected luciferase encoding mRNAs transcribed *in vitro* with either UTP or m^1^ΨTP into HEK293 cells, where they were translated. The base composition of the unmodified and modified mRNAs was assessed by liquid chromatography-mass spectrometry (LC-MS), and is consistent between the unmodified and modified mRNA species we generated (S1 Figure 8). We observed increased levels of luciferase protein expression in the m^1^Ψ-substituted mRNAs, consistent with previous reports (S1 Figure 9-10) (30). The luciferase proteins generated from both unsubstituted and m^1^Ψ-substituted mRNAs were purified and analyzed by mass spectrometry to identify amino acid substitutions.

Our mass spectrometry data analyses focused on a specific luciferase peptide with favorable ionization characteristics (Figure 2B) (30). ~1% of the amino acids in this peptide were substituted. This is a >20-fold increase over the level of amino acid substitution we previously observed for peptides generated from an unmodified version of the same luciferase reporter (− 0.05% of their amino acids substituted) (30). m^1^Ψ-mediated substitutions were detected on variety of codons (*e.g*. UUU, UAU), though we did not observe amino acid substitutions above background levels for every U-containing codon (*e.g*. UGG) (S1 Table 6). The highest levels of substitution were observed on the two Phe codons (UUU and UUC). Similar to our *in vitro* observations, serine and isoleucine/leucine amino acid substitutions were seen on both Phe codons with a > 6-fold increased frequency of substitution over peptide from unmodified mRNA (Figure 2B, S1 Tables S6-8) (30). Isoleucine and leucine have the same mass and therefore cannot be distinguished in this assay. We also noted that the likelihood of substitutions occurring was not uniform across m^1^Ψ containing codons. The levels of miscoding that we detect are consistent with what we would predict from our *in vitro* studies, as is the heterogeneity of amino acid substitution on m^1^Ψ-modified codons (Figures 2–3). Furthermore, the lack of uniformity in amino acid substitution was also seen in our previous findings indicating that Ψ also increases the levels of amino acid misincorporation in the same luciferase reporter peptide (30). Our results collectively suggest that the extent of misincorporation on any codon containing a C5-glyocside uridine isomer strongly depends on the sequence context in which the modification is present.

### Ψ-derived modifications change the energetics of mRNA:tRNA interactions in the ribosome A site

We sought to understand why Ψ and m^1^Ψ modifications alter the interactions between mRNAs and tRNAs during translation in a position dependent manner (Figures 1–4A). To approach this question we used molecular modeling (MM) and quantum mechanical calculations to examine unmodified and Ψ-, m^1^Ψ- and 3-methylpseudouridine (m^3^Y) modified UUU mRNA codons interacting with a cognate (tRNA^Phe(AAG)^) and near-cognate (tRNA^Ile(UAG)^) tRNAs in a portion of the ribosome A site (Figures 1A, 4). Although we did not investigate the translation of m^3^Ψ-containing codons, we included m^3^Ψ in our computational studies as a positive control for a modification that should severely perturb mRNA:tRNA interactions; methylation at the uridine N3 position removes the ability of uridine to donate a hydrogen bond and will limit tRNA binding. Our MM studies were conducted using models developed based on previously published crystal structure of the 70S *Thermus thermophilus* ribosome with tRNA^Phe^ bound on a ΨUU codon (30). The MM investigations were designed to examine how the location of a modification impacts the pairwise tRNA:mRNA interaction energies. Each modification was modeled in either the first, second, or third codon position and the energetics of tRNA^Phe/Ile^:mRNA interactions on modified codons were compared those on an unmodified UUU codon (ΔE) (Figure 4, S1 Figures 11 and 12). Only ΔE values with magnitudes ≳ 1 kcal/mol are considered large enough to potentially change mRNA:tRNA interactions.

We found that the incorporation of pseudouridine-derived modifications affect the energetic landscape for codon interactions with tRNA^Phe(GAA)^. As expected, the ΔE values for base pairing (ΔEbp) between each tRNA^Phe(GAA)^ anti-codon base and all three m^3^Ψ modified codons are significantly increased (ΔEbp = +8.6 to 19.7 kcal/mol, S1 Figure 11), suggesting the tRNA^Phe(GAA)^ is not likely to interact productively with m^3^Ψ-modified codon. Compared to m^3^Ψ, the effects of Ψ and m^1^Ψ that we observed on ΔE were not as uniform across the codon. ΨUU and UUΨ do not alter the energetics of base pairing (ΔE_bp,Ψ1:A36_ = −0.3 kcal/mol and ΔE_bp,Ψ3:G34_ = −0.4 kcal/mol), while substitutions in the second codon position (UΨU) are less energetically favorable (ΔE_bp,Ψ2:A35_ = +1.6 kcal/mol) (Figure 4AB). In contrast to Ψ, m^1^Ψ perturbs the ΔE_bp_ of codon:tRNA^Phe(GAA)^ anti-codon interactions when it is included at the first and third, but not second, positions of the codon. However, while ΔE_bp,m1Ψ1:A36_ is increased (+1.7 kcal/mol), this change is rendered insignificant by the compensatory alterations in energy between m^1^Ψ and the tRNA^Phe(GAA)^ bases (A35 and 2-methylthio-N6-isopentenyl-adenosine (ms^2^i^6^A)37) adjacent to the m^1^Ψ_1_:A36 interaction (ΔE_Σm1Ψ1:A35-ms2i6A37_= −1.4 kcal/mol) (Figure 4A-B). UUm^1^Ψ has more favorable energetics for interacting with tRNA^Phe(GAA)^ than UUU (ΔE_bp,m1Ψ3:G34_ = −2.9 kcal/mol), which are enhanced by m^1^Ψ interactions with the two adjacent tRNA nucleotides (ΔE_Σm1Ψ:U33-A35_ = −4.5 kcal/mol) (Figure 4B). These results are generally consistent with our kinetic studies demonstrating that the rate constant for Phe addition is modestly increased on UUm^1^Ψ codons relative to UUU, m^1^ΨUU and Um^1^ΨU (Figure 1B, S1 Table 1).

The substitution of U with Ψ, m^1^Ψ, and m^3^Ψ also impacts how codons interact with near cognate tRNA^Ile(GAU)^ in a manner supported by our *in vitro* translation kinetic findings (Figure 4A, C, D). Just as with tRNA^Phe(GAA)^, the ΔEbp values for individual m^3^Ψ-containing codon:tRNA^Ile(GAU)^ anticodon base pairs were typically less favorable relative to an unmodified UUU codon (up to +16.2 kcal/mol; S1 Figure 12). ΨUU and UΨU do not markedly influence ΔE_bp,Ψ1:U36_ and ΔE_bp,Ψ2:A35_ (+0.9 and +0.05 kcal/mol, respectively), though the energetics of tRNA^Ile(GAU)^ interactions with these substitutions are altered by ~1 kcal/mol when the ΔE values for the flanking tRNA nucleotides are also considered (ΔE_ΣΨ1:G34-U36_ = +1.1 kcal/mol, ΔE_ΣΨ2:A35-t6A37_= −0.8 kcal/mol) (Figure 4C). This differs from the effects of including Ψ at the third codon position. In the context of a UUΨ codon, ΔE_bp,Ψ3:G34_ is more favorable (−1.2 kcal/mol), but this effect is mitigated by the energetically less favorable interactions between the wobble Ψ and tRNA^Ile(GAU)^ nearby tRNA residues (ΔE_ΣU33-A35_ = −0.2 kcal/mol). Our findings support previous studies demonstrating that tRNA nucleotides adjacent to tRNA anticodon:codon base pairs are important determinants of mRNA decoding (62).

The trend for how m^1^Ψ substitutions alter codon:tRNA^Ile(GAU)^ anticodon base pairing interactions is similar to that of Ψ, however the magnitude of the changes is increased. For example, ΔE_bp,m1Ψ1:U36_ and ΔE_bp,m1Ψ3:G34_ are more negative (−1.6 kcal/mol and −3.4 and, respectively), while ΔE_bp,m1Ψ2:A35_ is largely unchanged (+0.3 kcal/mol) (Figure 4C). As we observed for Ψ, the extent of the m^1^Ψ-mediated changes to the codon:tRNA^Ile(GAU)^ anticodon interactions increases when ΔE is considered in the context of the flanking tRNA nucleotides (*e.g*. ΔE_Σm1Ψ1A35-t6A37_ = −9.2 kcal/mol). Our computational analyses generally support our experimental findings that Ψ-derived modifications differentially affect the interactions between codons and both cognate and near-cognate tRNAs in context dependent manner to alter mRNA:tRNA interactions in the ribosome decoding center (63–68) (Figure 4, S1 Figures 11-12).

## DISCUSSION

During the selection of aminoacylated-tRNAs the ribosome must compromise between the speed and accuracy of decoding. Chemical modification of the RNAs involved in decoding (*e.g*. mRNA and tRNA) permit the fine tuning of this balancing act. m^1^Ψ modifications are heavily used in mRNA-based therapeutics and vaccines and we were interested in determining how their inclusion in mRNA transcripts can impact translation elongation (33, 37). Our studies indicate that, depending on where it was located within a codon, m^1^Ψ either has no effect or modestly increases the rate constant (*k*_obs_) for cognate amino acid incorporation (Figure 1B). The rate constants (*k*_hyd,max_) for translation termination are not perturbed when sufficient concentrations of release factors are present (Figures 1C-D). Notably, the effect (if any) of m^1^Ψ on the ribosome reacting with near cognate tRNAs depending largely on the position of the modification within a codon (Figures 2-3). Our computational studies reveal that the context dependent effects of m^1^Ψ we observe might be explained by changes in the energetics of base pairing interactions between m^1^Ψ-substituted UUU codons and their cognate tRNA^Phe(GAA)^ (Figure 4AB), which vary depending on where m^1^Ψ is incorporated in the codon. Perturbations in base pairing energies have the ability to influence central steps in the translation elongation pathway including tRNA selection and accommodation. Although faster elongation rates might help to partially explain greater protein yield from m^1^Ψ containing transcripts in cells (S1 Figures 9-10), the preponderance of the increased yields in protein are likely due to m^1^Ψ-induced enhancements in mRNA stability and avoidance of the cell’s innate immune system (40, 41, 56, 69).

We found that m^1^Ψ alters not only cognate tRNA interactions with the ribosome, but also those of near-cognate tRNAs. Our kinetic investigations reveal that both m^1^Ψ and Ψ modifications can either enhance or limit amino acid substitution depending on aa-tRNA identity and the position of the modification within the codon (Figures 3 and 4A). These *in vitro* observations are supported by cellular reporter studies indicating that m^1^Ψ can promote miscoding events when included in full-length transcripts expressed in human cells (Figure 2B, S1 Tables 6, 8). The mass spectrometry assay implemented here does not have sufficient sensitivity to identify modified codons that exhibit reduced levels of amino acid substitution, as we might expect on Leu codons based on our kinetics. Our explorations of amino acid substitution in a peptide translated from a m^1^Ψ substituted mRNA are consistent with the increase in miscoding we previously observed on Ψ-containing mRNAs in the same experimental system (30). Comparison of miscoding rates on m^1^Ψ and Ψ-containing codons suggests that the addition of a methyl group, and altered ring electronics resulting from the exchange of the nitrogen, play distinct positional and codon specific roles in the m^1^Ψ-mediated modulation of miscoding (Figure 3). This is further supported by our MM calculations revealing that methylations (m^1^Ψ and m^3^Ψ) have larger impact on the energetics of tRNA:mRNA interactions than isomerization at the C5-position alone (Ψ) (Figure 4). These findings are in line with previous work demonstrating that naturally occurring mRNA modifications can differentially affect translation depending on their location within a codon or mRNA sequence (30, 31, 70).

In this work we go beyond observing that the sequence context of a modification matters for codon decoding (Figures 3 and 4) to try and identify *how* pseudouridine derived modifications change the fundamental interactions between mRNAs and tRNAs in a context dependent manner (Figure 4). Comparison of the rate constants for amino acid misincorporation on m^1^Ψ and Ψ containing codons, coupled with MM calculations provide evidence that fundamental changes in the energetics of mRNA:tRNA interactions may largely account for the context dependent outcomes we observe. Indeed, we find that the most energetically favorable interactions between the near cognate tRNA^Ile^ anticodon and Ψ and m^1^Ψ modified UUU Phe codons occur when these modifications are in the first and third position of the codon (Figure 4D-E). This is consistent with our kinetic studies indicating that Ile-tRNA^Ile^ reacts more rapidly with codons containing Ψ and m^1^Ψ in the first and third positions of a codon than in the second position (Figures 3, 4A). Additionally, our data suggest that tRNA ASL hypermodifications may also contribute to the context dependence of near-cognate decoding on modified Ψ and m^1^Ψ codons. We find that the rate constants for near cognate tRNAs possessing hypermodifications (t^6^A37 in tRNA^Ile(GAU)^, ms^2^i^6^A37 and cmo≡U34 in tRNA^Ser(UGA)^) are more sensitive to the position of mRNA modifications within a codon than the near cognate tRNA^Leu(CAG)^ that contains only a methylation at position 37 (m^1^G37) (Figure 4A). In line with this, MM calculations reveal that the hypermodifications adjacent to tRNA^Phe(GAA)^ and tRNA^Ile(GAU)^ anticodons strongly influence the energetics of codon:anticodon interactions with Ψ and m^1^Ψ, and can make tRNA interactions with a near-cognate codon up to 7.6 kcal/mol more energetically favorable. This correlates with our kinetic studies demonstrating that tRNA^Ile(GAU)^ reacts more rapidly on m^1^ΨUU, and is further supported by previous studies demonstrating that modified tRNA A37 nucleosides (t^6^A, ms^2^t^6^A, ct^6^A) improve the stability of the codon:anticodon duplex through enhanced base stacking (30, 41, 56, 57). Taken together, our kinetic and MM results suggest that both tRNA and mRNA modifications modulate the energetics of codon:anticodon interactions differently in varied sequence contexts to influence the decoding of Ψ and m^1^Ψ modified codons (Figure 4D,E and S1 Figure 7).

The ability of m^1^Ψ and Ψ to change the decoding behavior of the ribosome while only modestly altering amino acid addition and termination has several implications. Given the emerging evidence that Ψ is commonly included into mRNA at increased levels under cellular stress conditions, these findings support the possibility that Ψ-derived modifications can provide the cells with a way to transiently increase the diversity of the proteome under stress to increase fitness (15, 30, 73, 74). Furthermore, this could potentially be advantageous for mRNA vaccines relative to traditional vaccine platforms; greater antigen diversity might provide broader protection against circulating virus populations than single-strain vaccine formulations, similar to the increased efficacy of multivalent vaccines. While potentially beneficial in the context of mRNA vaccines, even small changes translational fidelity need to more carefully considered during the development of other classes of mRNA therapeutics. The complex rules governing the translational outcome of mRNA modifications such as m^1^Ψ are only beginning to be elucidated, and may in some cases prove to be critical to designing safe and effective mRNA therapeutics.

## ASSOCIATED CONTENT

### Supporting Information

Supporting Figures and Supporting Tables are available and associated with this text.

## AUTHOR INFORMATION

### Author Contributions

J.G. Monroe designed, performed and analyzed *in vitro* translation experiments, and participated in analyzing the MM data generated here. L. Mitchell and B. Roy designed and performed the mass spectrometry analyses of luciferase peptides translated in 293HEK cells. I. Deb and A. Frank designed, performed and the MM experiments. K.S. Koutmou designed and analyzed *in vitro* translation experiments, and participated in analyzing the MM data generated here. All authors contributed to writing the manuscript.

## ACKNOWLEDGMENT

We would like to acknowledge Dr. Dan Eyler and Tyler Smith for their thoughtful discussions and careful reading of the manuscript. We thank the following funding sources for their support: National Institutes of Health (NIGMS R35 GM128836), National Science Foundation (NSF CAREER 2045562 to K.S.K.), Research Corporation for Science Advancement (Cottrell Scholar Award to K.S.K.) and New England Biolabs (NEB) Inc. (M.Z.W. and B.R. are New England Biolabs, Inc. employees).

## ABBREVIATIONS

Phe, F: Phenylalanine
Ile, I: Isoleucine
Leu, L: Leucine
Ser, S: Serine
Ψ: Pseudouridine
m^1^Ψ: N1-methylpseudouridine
m^3^Ψ: N3-methylpseudouridine
ms^2^i^6^A: 2-methyl-thio-N6-isopentenyladenosine
t^6^A: N6-threonylcar-bamoyladenosine
ct^6^A: cyclic N6-threonylcar-bamoyladenosine
m^1^G: l-methylguanosine
cmo^5^U: uridine 5-oxyacetic acid

