## Supplemental Figures for "N1-Methylpseudouridine and pseudouridine modifications modulate mRNA decoding during translation"

**Dipeptide formation on an unmodified and $m^1\Psi$ modified UUU Phe codon**

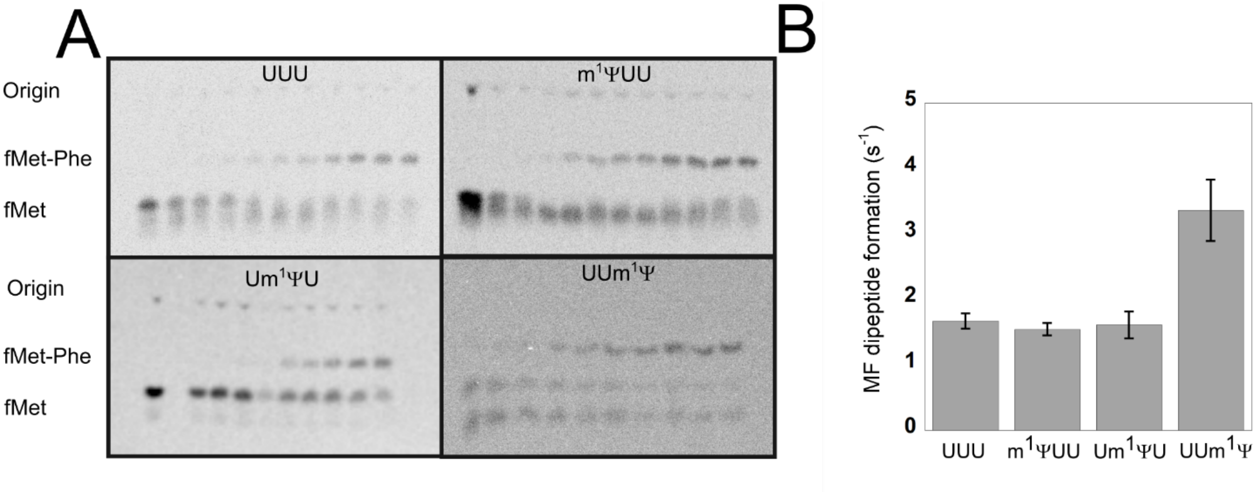


**Supplemental Figure 1. A.** Comparison of rates of dipeptide formation on a positionally $m^1\Psi$ modified Phe codon (UUU). **B.** N-formyl-[ 35S]- methionine-labeled peptides are separated by electrophoresis on a cellulose TLC in a volatile, acidic buffer, and detected by phosphorimaging. Representative images of of dipeptide formation acid addition assays, displaying the UUU control, $m^1\Psi$UU Um$^1\Psi$U UUm$^1\Psi$ are shown. The brightness and contrast of these images have been adjusted to clearly show all bands and the background, and as a consequence pixel intensity is no longer linear with signal.

**Time courses of peptide hydrolysis by Release Factor 1 on an unmodified and $m^1\Psi$ modified UAA stop codon**

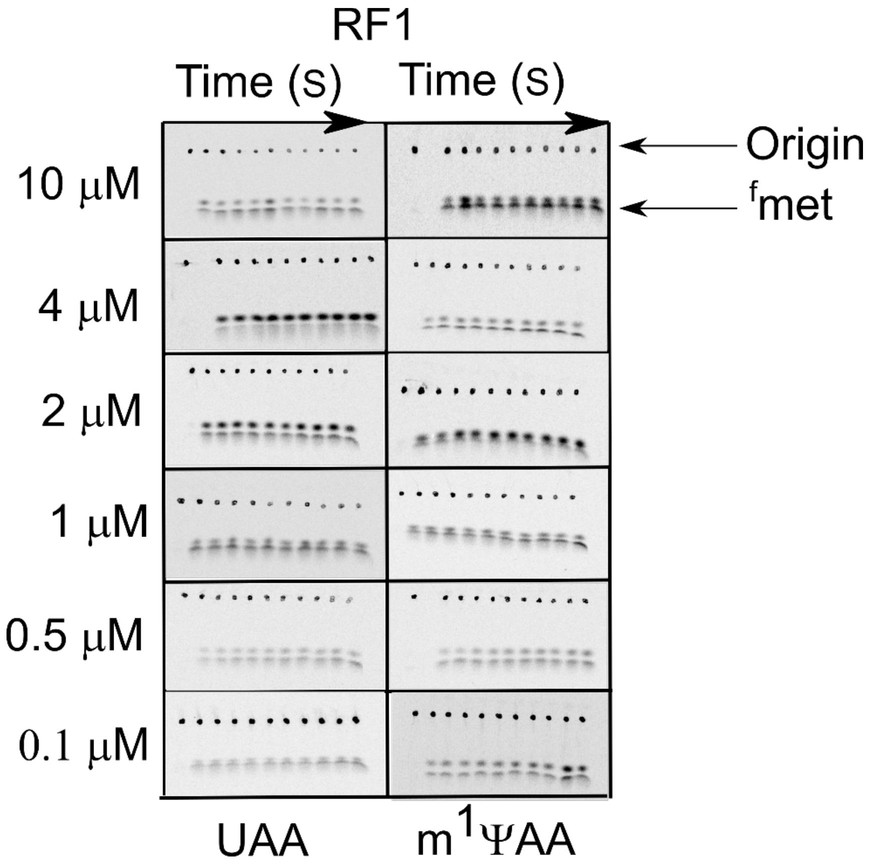


**Supplemental Figure 2.** Representative images of time courses for release factor 1 are shown. The brightness and contrast of these images have been adjusted to clearly show all bands and the background, and as a consequence pixel intensity is no longer linear with signal.

**Time courses of peptide hydrolysis by Release Factor 2 on an unmodified and $m^1\Psi$ modified UAA stop codon**

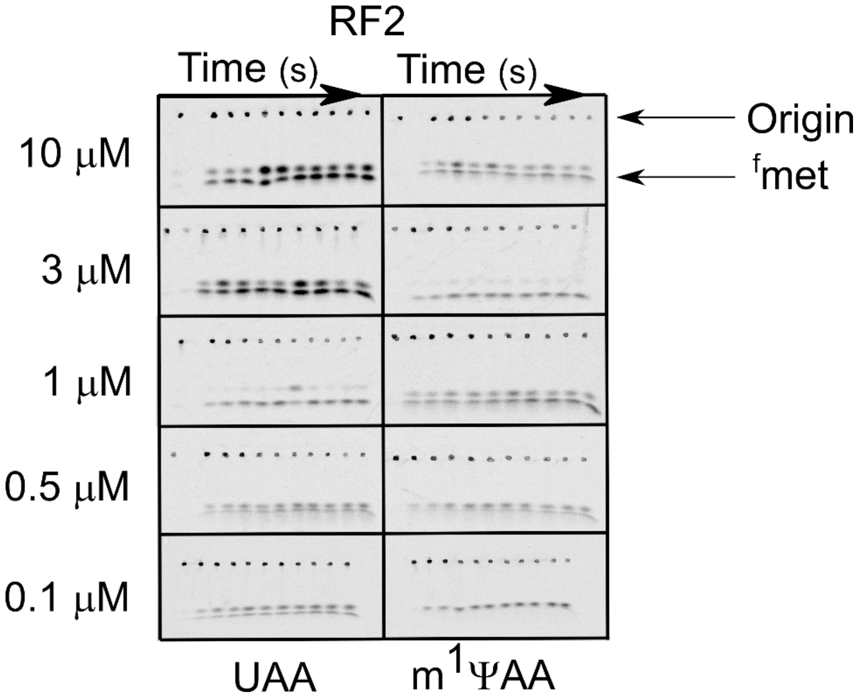


**Supplemental Figure 3.** Representative images of time courses for release factor 2 are shown. The brightness and contrast of these images have been adjusted to clearly show all bands and the background, and as a consequence pixel intensity is no longer linear with signal.

**End point screening assays for changes in miscoding levels on m$^1\Psi$ modified UUU codons**

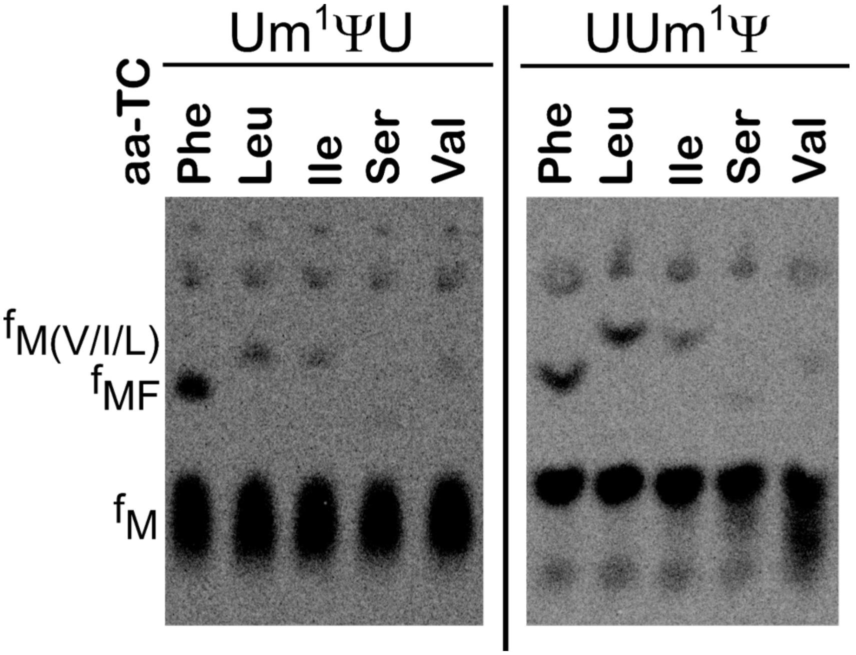


**Supplemental Figure 4.** The endpoint mis-incorporation defects of a m$^1\Psi$ positioned at the second or third positions of a Phe UUU codon. The IC was reacted with total tRNA where only one species of tRNA was charged. The miscoded dipeptide were separated via eTLC and the miscoded products identified by size and charge.

**Titration of Ile tRNA on a phenylalanine codon**

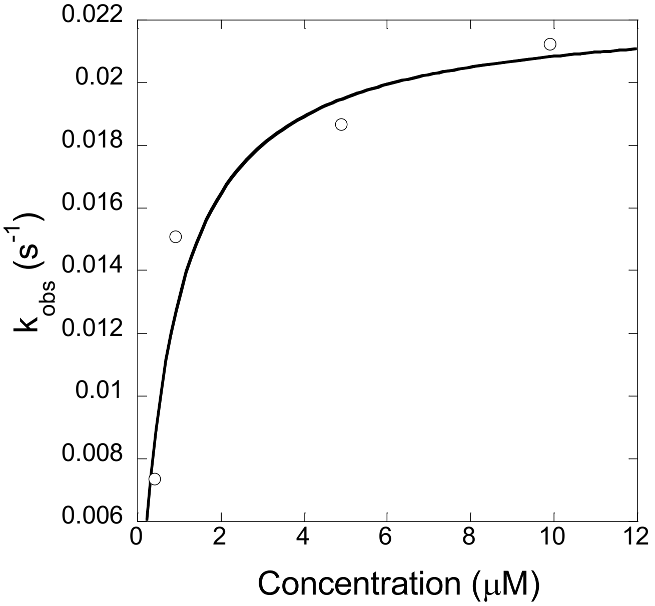


**Supplemental Figure 5.** In order to ensure that the rates of miscoding reactions were carried out under saturating conditions the $K_{1/2}$ of Ile miscoding by $tRNA^{Ile(GAU)}$ was determined. A $K_{1/2}$ of approximately 0.7 μM was determined, so a final of concentration of 10 μM aa-tRNA to ensure that the miscoding assays were conducted under saturating conditions.

**Representative Images of Misincorporated Dipeptide Products an Unmodified and $m^1\Psi$ Modified Phenylalanine Codon**

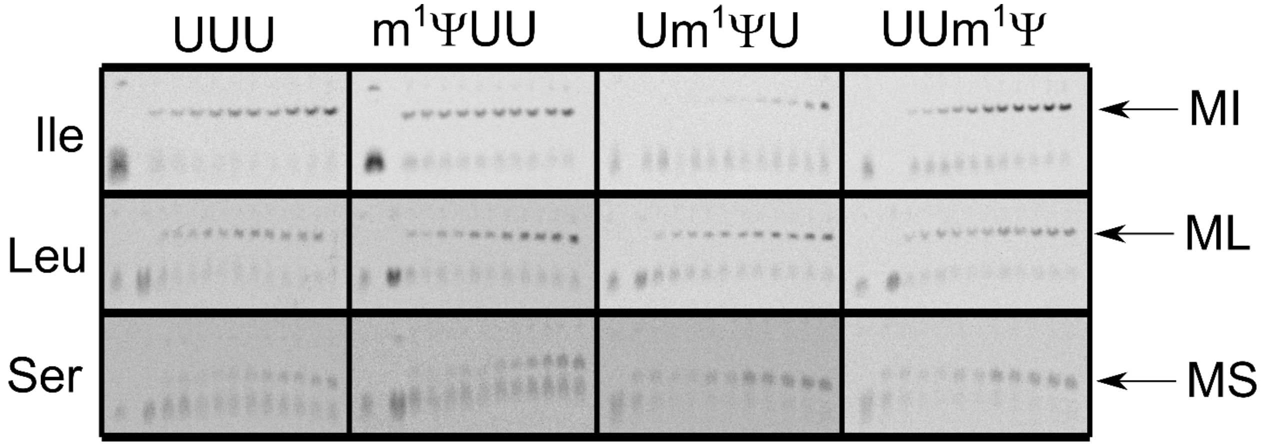


**Supplemental Figure 6.** Representative eTLCs images of of dipeptide formation acid addition assays, displaying the UUU control, $m^1\Psi$UU U$m^1\Psi$U UU$m^1\Psi$ are shown.of the miscoding assays performed. Note the reduced formation of the MI dipeptide when the second position is modified with $m^1\Psi$.

**Representative Images of Misincorporated Dipeptide Products an Unmodified and Ψ Modified Phenylalanine Codon**

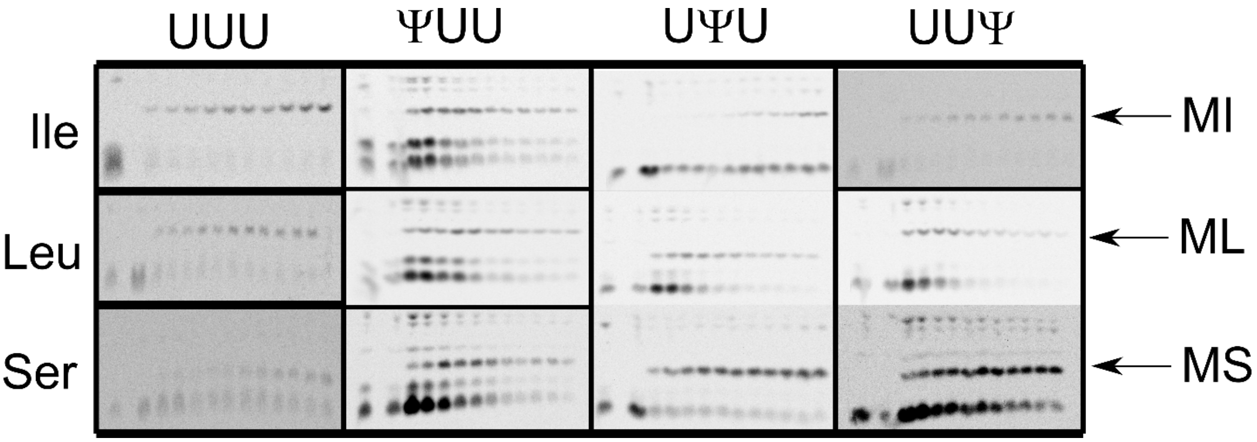


**Supplemental Figure 7.** Representative eTLCs images of of dipeptide formation acid addition assays, displaying the UUU control, m$^{1}$ΨUU Um$^{1}$ΨU UUm$^{1}$Ψ are shown.of the miscoding assays performed. Note the reduced formation of the MI dipeptide when the second position is modified with Ψ.

### Mass Spectrum Analysis of RNA Bases and Incorporation Efficiency

A

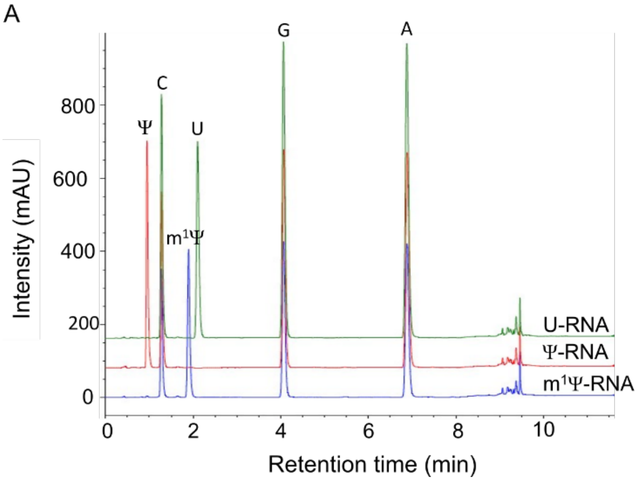


B

| Nucleotide | U | Ψ | m[1]Ψ |
|---|---|---|---|
| rA | 0.92 | 0.91 | 0.93 |
| rC | 0.83 | 0.85 | 0.83 |
| rG | 1 | 1 | 1 |
| rU/Ψ/m[1]Ψ | 0.71 | 0.73 | 0.72 |

**Supplemental Figure 8.** (A) Base composition of each RNA base detected by LC-MS, relative to guanosine. (B) Incorporation efficiency of modified nucleotides was assessed by Liquid Chromatography-Mass Spectrometry (LC-MS).

**Expression of Unmodified and Modified Luciferase mRNA in HEK293 Cells**

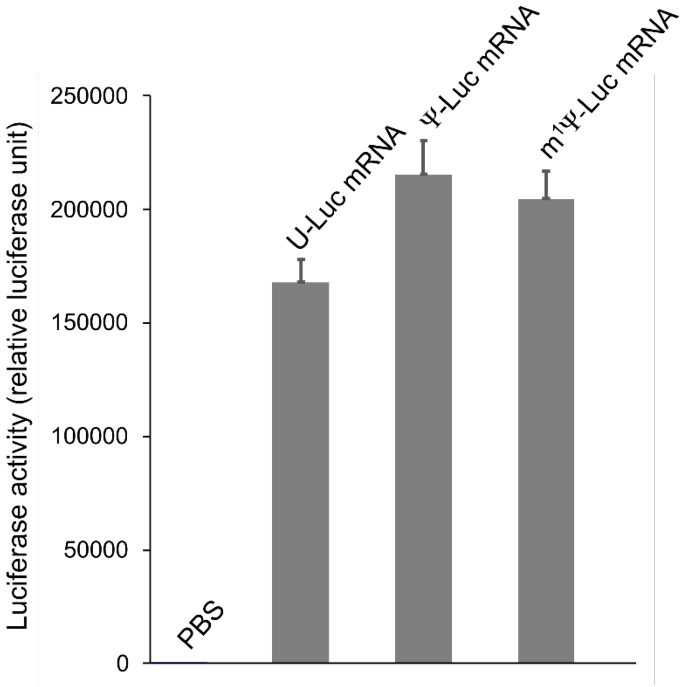


**Supplemental Figure 9.** Expression of m$^1\Psi$ modified luciferase mRNA *in vivo*. Luciferase activity assay demonstrating functionality of unmodified and modified luciferase mRNAs.

### Protein Yields from Unmodified and Modified Luciferase mRNA in HEK293 Cells

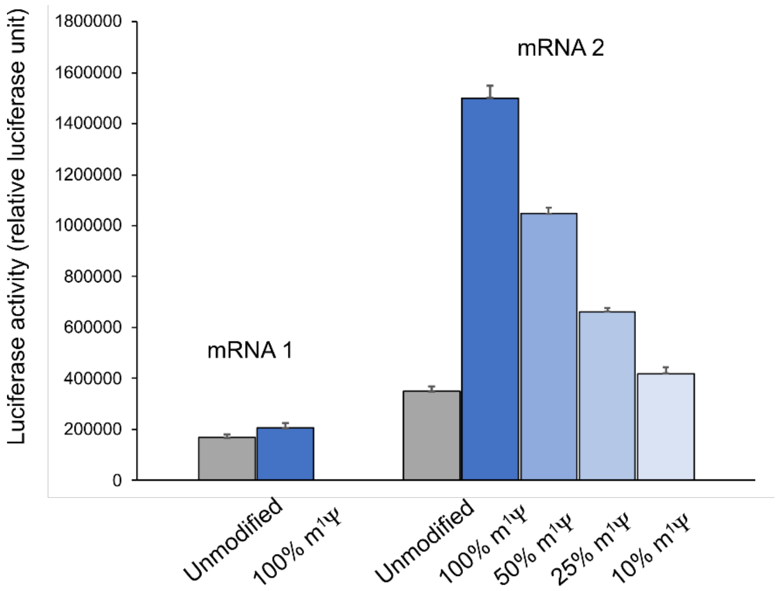


**Supplemental Figure 10.** The yield of active protein from $m^1\Psi$ modified mRNAs depends on the level of modification and sequence context. Two mRNAs coding for luciferase but with silent coding region changes were in vitro transcribed in the presence of varying ratios of uridine and $m^1\Psi$triphosphate. Purified mRNAs were transfected into cultured mammalian cells and luciferase activity was assayed at a single time point.

### Change in energy based on Molecular Modeling of cognate decoding by tRNA$^{Phe}$

| Interacting Nucleotide | Phe tRNA with the MIA37 ΔE | | | | | | | | |
|---|---|---|---|---|---|---|---|---|---|
| | | Ψ | | | m1Ψ | | | m3Ψ | |
| | **1st** | **2nd** | **3rd** | **1st** | **2nd** | **3rd** | **1st** | **2nd** | **3rd** |
| ADE16 | -0.487 | 0.015 | 0.375 | -0.658 | -0.789 | -0.702 | 0.017 | 0.461 | 0.244 |
| URA17 | -0.547 | 0.048 | 0.25 | -0.836 | -1.091 | -1.094 | 0.019 | 0.677 | 0.28 |
| GUA18 | -6.168 | -0.646 | 0.319 | -5.851 | -3.572 | -2.207 | -5.225 | 0.536 | 0.387 |
| URA19/URA20 | -1.716 | -0.643 | 0.517 | -1.784 | -6.604 | -7.131 | -0.069 | -2.238 | 0.269 |
| URA19/URA21 | -0.495 | -0.09 | 0.693 | -0.729 | -3.758 | -6.05 | 0.135 | 1.79 | 1.317 |
| URA22 | -0.465 | 0.039 | 2.038 | -0.528 | -1.082 | -0.287 | -0.083 | 0.755 | 2.204 |
| ADE23 | 0.034 | 0.016 | -0.051 | 0.007 | 0.074 | 0.13 | -0.009 | -0.009 | -0.18 |
| GUA28 | -0.35 | 0.06 | 0.31 | -0.406 | -0.626 | -0.498 | 0.083 | 0.6 | 0.247 |
| GUA29 | -0.343 | 0.058 | 0.328 | -0.417 | -0.651 | -0.493 | 0.076 | 0.596 | 0.24 |
| GUA30 | -0.289 | 0.079 | 0.33 | -0.351 | -0.59 | -0.438 | 0.112 | 0.601 | 0.253 |
| ADE31 | -0.468 | -0.008 | 0.306 | -0.569 | -0.753 | -0.519 | -0.103 | 0.538 | 0.247 |
| PSU32 | -0.503 | 0.066 | 0.375 | -0.622 | -0.893 | -0.64 | 0.07 | 0.797 | 0.344 |
| URA33 | -0.524 | 0.086 | 0.4 | -0.665 | -1.13 | -0.855 | -0.155 | 0.961 | 0.445 |
| **GUA34** | -0.82 | -0.251 | -0.417 | -1.002 | -1.79 | -2.939 | -0.211 | -0.278 | 19.673 |
| **ADE35** | -0.37 | 1.631 | 0.47 | -0.443 | 0.45 | -0.669 | 0.472 | 14.527 | -0.327 |
| **ADE36** | -0.295 | -0.277 | 0.305 | 1.742 | -2.534 | -1.014 | 8.56 | 0.27 | 0.427 |
| MIA37 | -0.542 | 0.162 | 0.403 | -1.358 | -0.939 | -0.913 | -1.341 | 1.05 | 0.361 |
| ADE38 | -0.541 | 0.085 | 0.332 | -0.727 | -0.87 | -0.728 | 0.177 | 0.763 | 0.309 |
| PSU39 | -0.436 | 0.062 | 0.297 | -0.601 | -0.68 | -0.512 | 0.086 | 0.555 | 0.251 |
| CYT40 | -0.607 | -0.059 | 0.204 | -0.743 | -0.739 | -0.524 | -0.273 | 0.367 | 0.172 |
| CYT41 | -0.358 | 0.002 | 0.197 | -0.442 | -0.52 | -0.394 | -0.091 | 0.343 | 0.16 |
| CYT42 | -0.028 | -0.007 | -0.004 | -0.015 | 0.01 | 0.02 | -0.05 | -0.022 | 0.001 |

| | |
|---|---|
| (yellow) | mRNA |
| (green) | tRNA |

**Supplemental Figure 11.** The change in energy (ΔE kcal/mol) for each codon position interacting with nearby mRNA residues (A16-A23) and tRNA$^{Phe(GAA)}$ positions G28 to C42 on a Ψ, m$^1$Ψ and m$^3$Ψ modified codon relative to unmodified UUU codon. The values used to discern $\Delta E_{bp}$ and $\Delta E_{\Sigma\Psi/\mu1\Psi:tRNApositionsx\text{-}y}$ in the text are highlighted in the dotted box.

### Change in energy based on Molecular Modeling of cognate decoding by $tRNA^{Ile}$

| | Ile tRNA ΔE | | | | | | | | |
|---|---|---|---|---|---|---|---|---|---|
| | Ψ | | | m1Ψ | | | m3Ψ | | |
| Interacting Nucleotide | 1st | 2nd | 3rd | 1st | 2nd | 3rd | 1st | 2nd | 3rd |
| ADE16 | -0.179 | 0.034 | 0.263 | -0.977 | -0.66 | -0.709 | 0.046 | 0.484 | 0.099 |
| URA17 | -0.128 | 0.059 | 0.299 | -1.302 | -0.943 | -1.122 | 0.239 | 0.703 | 0.051 |
| GUA18 | -3.772 | -0.626 | 0.281 | -6.715 | -3.299 | -2.243 | -2.626 | 0.561 | -0.126 |
| URA19 | -0.745 | -0.631 | 0.181 | -2.678 | -5.758 | -7.184 | 0.778 | -2.262 | -1.041 |
| URA19 | -0.185 | 0.247 | 0.432 | -1.222 | -3.517 | -6.091 | -0.172 | 4.289 | 1.902 |
| URA22 | -0.186 | 0.088 | 1.12 | -0.931 | -0.941 | -0.683 | -0.07 | 0.668 | 0.874 |
| ADE23 | -0.003 | 0.004 | -0.093 | 0.002 | 0.053 | 0.063 | -0.019 | -0.049 | -0.357 |
| ADE28 | -0.131 | 0.089 | 0.29 | -0.71 | -0.452 | -0.414 | 0.001 | 0.549 | 0.267 |
| CYT29 | -0.163 | 0.052 | 0.281 | -0.774 | -0.517 | -0.442 | -0.108 | 0.501 | 0.194 |
| CYT30 | -0.088 | 0.044 | 0.266 | -0.696 | -0.458 | -0.376 | 0.021 | 0.512 | 0.213 |
| CYT31 | -0.162 | 0.037 | 0.253 | -0.881 | -0.542 | -0.462 | -0.046 | 0.514 | 0.221 |
| CYT32 | -0.316 | -0.008 | 0.284 | -1.341 | -0.781 | -0.625 | -0.082 | 0.636 | 0.262 |
| URA33 | -0.249 | 0.029 | 0.358 | -1.414 | -0.993 | -0.832 | -0.035 | 0.948 | 0.325 |
| **GUA34** | -0.528 | -0.174 | -1.162 | -1.602 | -1.363 | -3.364 | -0.438 | 0.707 | 14.778 |
| **ADE35** | -0.157 | 0.871 | 0.6 | -0.842 | 0.304 | -0.469 | 0.003 | 16.205 | 1.309 |
| **URA36** | 0.05 | 0.357 | 0.448 | -1.594 | -0.741 | -0.85 | -3.427 | -0.379 | 0.287 |
| T6A37 | -0.664 | 0.202 | 0.775 | -6.803 | -2.005 | -1.713 | -1.202 | 1.781 | 0.481 |
| ADE38 | -0.029 | 0.177 | 0.364 | -1.688 | -0.717 | -0.726 | 0.229 | 0.857 | 0.184 |
| GUA39 | -0.31 | 0.017 | 0.255 | -1.188 | -0.677 | -0.58 | -0.101 | 0.517 | 0.148 |
| GUA40 | -0.226 | 0.019 | 0.236 | -1.048 | -0.583 | -0.52 | -0.009 | 0.527 | 0.162 |
| GUA41 | -0.2 | 0.018 | 0.191 | -0.807 | -0.459 | -0.431 | -0.087 | 0.395 | 0.137 |
| URA42 | -0.008 | -0.016 | 0.002 | -0.022 | -0.023 | -0.009 | 0.006 | 0.018 | -0.009 |

| | |
|---|---|
| (yellow) | mRNA |
| (green) | tRNA |

**Supplemental Figure 12.** The change in energy (ΔE kcal/mol) for each codon position interacting with nearby mRNA residues (A16-A23) and $tRNA^{Ile(GAU)}$ positionss A28 to U42 on a Ψ, $m^1Ψ$ and $m^3Ψ$ modified codon relative to unmodified UUU codon. The values used to discern $ΔE_{bp}$ and $ΔE_{ΣΨ/μ1Ψ:tRNApositionsx-y}$ in the text are highlighted in the dotted box.

**$m^1\Psi$ Modified mRNA sequences prepared by Dharmacon.**

1st Position Modified:
5'G.G.U.G.U.C.U.U.G.C.G.A.G.G.A.U.A.A.G.U.G.C.A.U.U.A.U.G.$m^1\Psi$.U.U.U.A.A.G.C.C.C.U.U.C.U.G.U.A.G.C.C.A 3'

2nd Position Modified:
5'G.G.U.G.U.C.U.U.G.C.G.A.G.G.A.U.A.A.G.U.G.C.A.U.U.A.U.G.U.$m^1\Psi$.U.U.A.A.G.C.C.C.U.U.C.U.G.U.A.G.C.C.A 3'

Phe UUU Codon 3rd Position Modified:
5'G.G.U.G.U.C.U.U.G.C.G.A.G.G.A.U.A.A.G.U.G.C.A.U.U.A.U.G.U.U.$m^1\Psi$.U.A.A.G.C.C.C.U.U.C.U.G.U.A.G.C.C.A 3'

UAA Stop Codon 1st Position Modified:
5'G.G.U.G.U.C.U.U.G.C.G.A.G.G.A.U.A.A.G.U.G.C.A.U.U.A.U.G.$m^1\Psi$.A.A.G.U.U.G.C.C.U.U.C.U.G.U.A.G.C.C.A 3'

**Supplemental Figure 13.** Dharmacon prepared $m^1\Psi$ modified mRNA sequences for the Phe (UUU) codon at the 1st, 2nd, 3rd position and the first position of UAA stop codon. The transcripts were HPLC purified and converted to 2'-hydroxyl form and desalted.

### Quality Control Test of mRNA Sequences

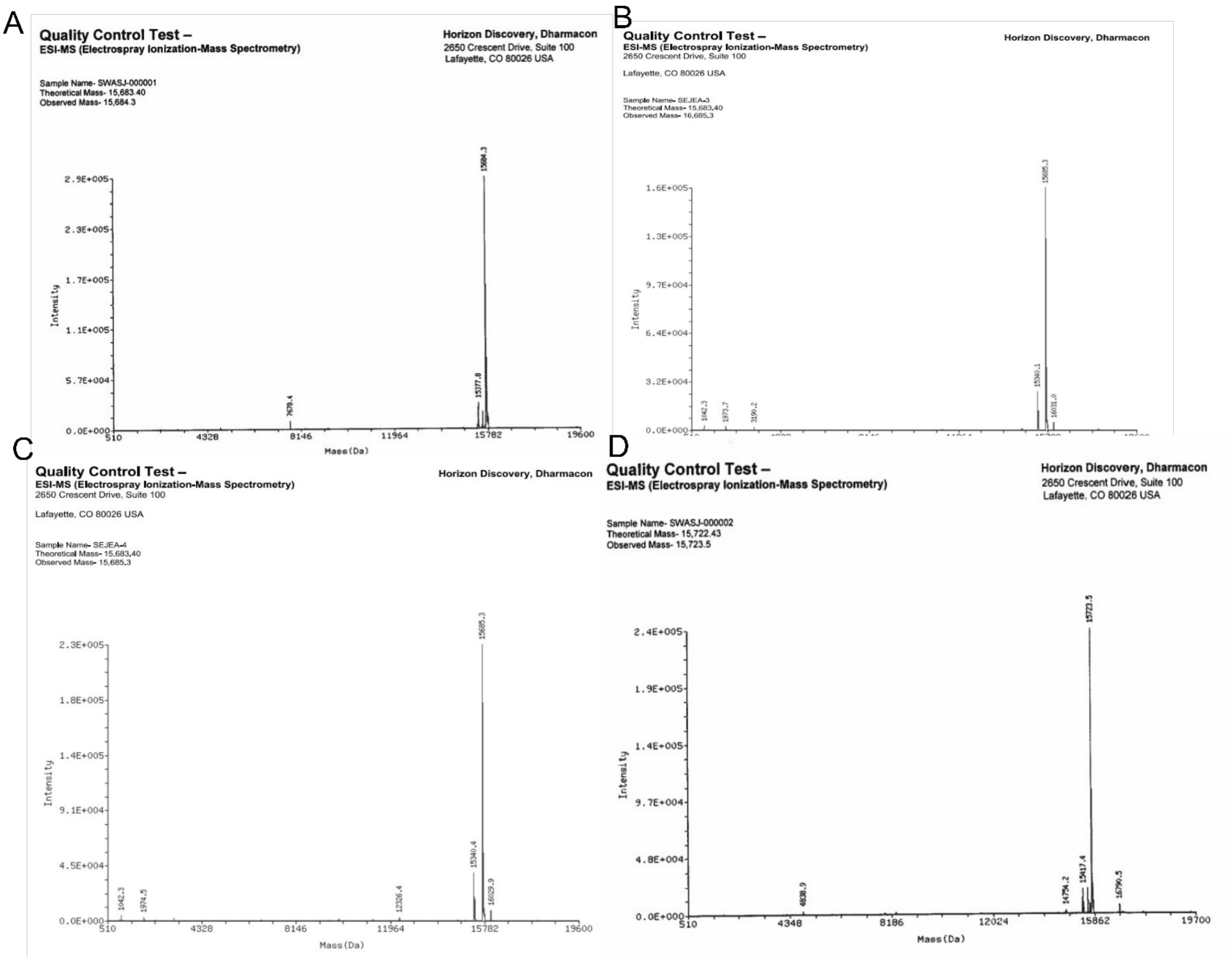


**Supplemental Figure 14.** Dharmacon quality control of $m^1\Psi$ modified mRNA sequences by ESI-MS (**A**) first position of Phe UUU modified to $m^1\Psi$UU (**B**) ) second position of Phe UUU modified to U$m^1\Psi$U, (**C**) third position of Phe UUU modified to UU$m^1\Psi$, (**D**) first position of a UAA stop codon modified to $m^1\Psi$AA.
