## Supplemental Tables for "N1-Methylpseudouridine and pseudouridine modifications modulate mRNA decoding during translation"

**Amino acid addition on an unmodified and m$^1\Psi$ modified phenylalanine codons**

| Rates of dipeptide formation | | |
|---|---|---|
| **Codon** | **Rate ($s^{-1}$)** | **Error** |
| UUU | 1.7 | 0.1 |
| m$^1\Psi$UU | 1.5 | 0.1 |
| Um$^1\Psi$U | 1.6 | 0.2 |
| UUm$^1\Psi$ | 3.4 | 0.5 |

**Supplemental Table 1.** This table reflects the values derived from plots in Figure 1B displaying the formation of dipeptide (MetPhe) catalyzed by the ribosome on unmodified and modified UUU codons. The reported $k_{obs}$ and standard error values are from the fit of a single curve to all replicate time courses.

**Titration of Release Factor 1 on an unmodified and $m^1\Psi$ modified UAA stop codon**

| Release Factor 1 Assays ($k_{hyd}$ $s^{-1}$) | | | | |
|---|---|---|---|---|
| **Concentration (μM)** | **UAA** | **Error** | **$m^1\Psi$AA** | **Error** |
| **0.1** | **0.06** | **0.022** | **0.01** | **0.002** |
| **0.5** | **0.04** | **0.009** | **0.02** | **0.003** |
| **1.0** | **0.10** | **0.011** | **0.05** | **0.008** |
| **2.0** | **0.15** | **0.024** | **0.08** | **0.013** |
| **4.0** | **0.08** | **0.009** | **0.06** | **0.008** |
| **10.0** | **0.11** | **0.012** | **0.12** | **0.001** |

**Supplemental Table 2.** Pre-termination complexes were prepared on mRNAs containing the coding sequences AUG-UAA-GUU or AUG-$m^1\Psi$AA-GUU. Peptide release assays were performed in 1 X 219-Tris buffer at room temperature (70 nM pre-TCs, RF1 ranging from 100 nM to 10 μM final concentration). Reaction aliquots were quenched with 5% formic acid (final) at varying time points. Free f-[ $^{35}$S]-Met was separated from f-[ 35S]-Met-tRNA$^{fMet}$ by electrophoretic TLC and quantified by phosphorimaging.

**Titration of Release Factor 1 on an unmodified and $m^1\Psi$ modified UAA stop codon**

| Release Factor 2 Assays ($k_{hyd}$ $s^{-1}$) | | | | |
|---|---|---|---|---|
| **Concentration (μM)** | **UAA** | **Error** | **$m^1\Psi$AA** | **Error** |
| **0.1** | **0.01** | **0.003** | **0.003** | **0.005** |
| **0.5** | **0.02** | **0.004** | **0.02** | **0.005** |
| **1.0** | **0.05** | **0.009** | **0.03** | **0.005** |
| **3.0** | **0.05** | **0.005** | **0.04** | **0.004** |
| **10.0** | **0.06** | **0.006** | **0.05** | **0.006** |

**Supplemental Table 3.** Pre-termination complexes were prepared on mRNAs containing the coding sequence AUG-UAA-GUU or AUG- $m^1\Psi$AA-GUU. Peptide release assays were performed in 1X219-Tris buffer at room temperature (70 nM pre-TCs, RF2 ranging from 100 nM to 10 μM final concentration). Reaction aliquots were quenched with 5% formic acid (final) at varying time points. Free f-[ $^{35}$S]-Met was separated from f-[ 35S]-Met-tRNAfMet by electrophoretic TLC and quantified by phosphorimaging.

**Rates of misincorporation on an unmodified and $m^1\Psi$ modified Phe UUU codon**

| | Amino acid substitution $k_{obs}$ ($s^{-1}$) | | | | | |
|---|---|---|---|---|---|---|
| | **Ile** | **Error** | **Leu** | **Error** | **Ser** | **Error** |
| **UUU** | 0.018 | 0.0028 | 0.04 | 0.008 | 0.004 | 0.00017 |
| **$m^1\Psi$UU** | 0.04 | 0.003 | 0.03 | 0.003 | 0.003 | 0.0004 |
| **U$m^1\Psi$U** | 0.0018 | 0.00025 | 0.01 | 0.0015 | 0.014 | 0.0013 |
| **UU$m^1\Psi$** | 0.013 | 0.0009 | 0.019 | 0.003 | 0.0032 | 0.0004 |

**Supplemental Table 4**. The $k_{obs}$ for dipeptide formation on unmodified and $m^1\Psi$ positionally modified UUU codons. The reported $k_{obs}$ and standard error values are from the fit of a single curve to 3 replicate time courses normalized to an end-point of 100% (shown in Figure 3).

**Rates of misincorporation on an unmodified and Ψ modified UUU Phe codon**

| | Amino acid substitution $k_{obs}$ ($s^{-1}$) | | | | | |
|---|---|---|---|---|---|---|
| | **Ile** | **Error** | **Leu** | **Error** | **Ser** | **Error** |
| **UUU** | 0.018 | 0.0028 | 0.04 | 0.008 | 0.004 | 0.00017 |
| **ΨUU** | 0.018 | 0.0029 | 0.0180 | 0.0025 | 0.016 | 0.0014 |
| **UΨU** | 0.00175 | 0.00026 | 0.014 | 0.0005 | 0.016 | 0.0021 |
| **UUΨ** | 0.0073 | 0.00128 | 0.022 | 0.0015 | 0.0069 | 0.0004 |

**Supplemental Table 5.** The $k_{obs}$ for dipeptide formation on unmodified and Ψ positionally modified UUU codons. The reported $k_{obs}$ and standard error values are from the fit of a single curve to 3 replicate time courses normalized to an end-point of 100% (shown in Figure 3).

**Frequency of amino acid substitution observed on m$^1\Psi$ containing codons in 293HEK cells by mass spectrometry**

| | Substitutions observed | Frequency of substitution (%) |
|---|---|---|
| Phe | Ser, Ile | 0.31 |
| Tyr | Cys, His | 0.13 |
| Leu | Pro, Ser | 0.22 |
| Ile | Thr, Trp, Val | 0.11 |
| Val | Glu, Ala | 0.14 |
| Trp | None observed | 0 |

**Supplemental Table 6.** This table summarizes the amino acid substitutions detected in the U-containing codons in the entire luciferase dataset for multiple peptides when mRNAs were synthesized with $\Psi$. For calculating the frequencies, an extracted ion chromatogram was generated at < 5ppm for each of the peptides of interest from the total ion current, and the area under the curve for each EIC was calculated. This was then used to calculate the percentage of substitution (area under the curve for peptides with a specific substitution/[area under the curve for all wild-type peptides with no substitution + area under the curve for all peptides with substitutions]).

**$m^1\Psi$ containing codons in luciferase mRNA**

| Amino acids evaluated for miscoding in 293H cells | Codons evaluated for miscoding in 293H cells |
|---|---|
| Phe | UUU, UUC |
| Leu | UUA, UUG, CUU, CUC, CUA, CUG |
| Ile | AUU, AUC, AUA |
| Val | GUU, GUC, GUA, GUG |
| Trp | UGG |
| Tyr | UAU, UAC |

**Supplemental Table 7.** Uridine-containing codons analyzed for elongation miscoding.

**$m^1\Psi$ substituted codons and their corresponding miscoding events**

| Codon | Substitutions observed | Frequency of substitution (%) |
|---|---|---|
| Phe | Ser, Ile/Leu | 0.31% |
| Tyr | Cys, His | 0.14% |
| Leu | Pro, Ser | 0.36% |

**Supplemental Table 8.** This table summarizes the amino acid substitutions detected from U-containing Phe, Tyr, Leu codons in the KGPAPFYPLEDGTAGEQLHK peptide when mRNAs were synthesized with $m^1\Psi$. The frequencies of the substitutions are also denoted. For calculating the frequencies, an extracted ion chromatogram was generated at <5ppm for each of the peptides of interest from the total ion current, and the area under the curve for each EIC was calculated. This was then used to calculate the percentage of substitution (area under the curve for peptides with a specific substitution/[area under the curve for all wild-type peptides with no substitution + area under the curve for all peptides with substitutions]).
